# A method for simultaneous targeted mutagenesis of all nuclear rDNA repeats in *Saccharomyces cerevisiae* using CRISPR-Cas9

**DOI:** 10.1101/276220

**Authors:** Lilly Chiou, Daniele Armaleo

## Abstract

*Saccharomyces cerevisiae* has been the prime model to study the assembly and functionality of eukaryotic ribosomes. Within that vast landscape, the specific problem of mutagenizing all 150 nuclear rRNA genes was bypassed using strains whose chromosomal copies had been deleted and substituted by plasmid-borne rDNA. Work with these strains has produced important insights, but nucleolar structure is altered and such yeast-specific approaches are elaborate and not transferable to most other eukaryotes. We describe here a simple CRISPR-Cas9 based method to place targeted mutations in all 150 chromosomal rDNA repeats in yeast. The procedure *per se* is not expected to alter the nucleolus and is potentially applicable also to other eukaryotes. Yeast was transformed with a plasmid bearing the genes for Cas9 and for the guide RNA, engineered to target a site in the SSU region. Our mutagenesis plan included insertion of a spliceosomal intron in the normally intronless yeast nuclear rDNA. Despite the potential lethality of cutting all 150 rDNA repeats at the same time, yeast survived the Cas9 attack through inactivation of the cut sites either by point mutations or by inserting the intron, which was spliced out correctly from the rRNA transcript. In each mutant strain the same mutation was present in all rDNA repeats and was stably inherited even after removal of the Cas9 plasmid.

## Introduction

CRISPR-Cas9 (“CRISPR”) can simplify and improve genome engineering even in an already very powerful and versatile model for gene manipulation like *Saccharomyces cerevisiae* (8; 24). We describe here the use of CRISPR to mutagenize in a single step the entire set of rDNA repeats in *S. cerevisiae*. Mutagenesis of rDNA has been key in understanding the functionality of eukaryotic ribosomes using yeast as a model (6; 20; 23). However, due to the large numbers of yeast rDNA copies (∼150), those experiments involved the use of specially constructed strains lacking chromosomal rDNA but surviving with multi-copy plasmids bearing each one rRNA gene which could be mutagenized *in vitro* (29). A similar approach was used to modify the seven rRNA genes of *E. coli* (2). The insertion of the rDNA intron from *Tetrahymena thermophila* into yeast rDNA at its unique insertion site (16; 18), foreshadowed in the nineties some of the characteristics of CRISPR: as the intron endonuclease cut every wild type rDNA repeat, lethality could only be avoided if the cut site was modified by intron insertion or by mutation. Recently, a novel method was described (14) to extensively mutagenize the single 16S rDNA copy (*rrs*) in the synthetic 1.08 Mb genome of the bacterium *Mycoplasma mycoides* (10). CRISPR was used to introduce mutated *rrs* copies into *M. mycoides* genomes that had been transplanted into yeast. The modified genomes were then reintroduced into *M. mycoides*. Such a protocol might be difficult to adapt to most eukaryotes, with their large genomes and hundreds of nuclear rDNA genes. In this first direct application of CRISPR to eukaryotic rDNA mutagenesis, we targeted the 18S region in yeast as a proof of concept. Successful targeting of yeast rDNA with CRISPR requires the cell to withstand the concurrent cutting of ∼150 essential and tandemly repeated gene copies. Yeast survived, bearing either point mutations or even an intron, which are spread uniformly across all chromosomal rDNA repeats and stably inherited. The method is relatively simple and should also be transferable to other eukaryotes.

## Materials and Methods

### Plasmids, strains, and media

DH5alpha *E. coli* cells containing the plasmid pCAS were obtained from Addgene (#60847). Plasmid pCAS is a kanamycin/G418 shuttle vector carrying the gene for Cas9 and a generic guide RNA expression cassette (24). Plasmid pCAS-SS2-0 is the pCAS derivative we constructed to direct Cas9 to the SSU rDNA (see next section). Competent *E. coli* strain DH10B (New England Biolabs, C3019I) was used for transformations. To prepare competent *S. cerevisiae*, the protocol of Ryan et al. (25) was followed. Strain YJ0 (27) (*MAT****a***, *gal4*Δ, *gal80*Δ, *ura3–52, leu2–3, 112 his3, trp1, ade2–101*) was used, originally provided by Stephen Johnston. *E. coli* was grown in LB medium (doi:10.1101/pdb.rec8141) at 37°C. For transformant selection, kanamycin (ThermoFisher Scientific) was added to the LB medium to 100 mg/L. Yeast was grown in YEPD medium (doi:10.1101/pdb.rec12161) at 30°C. To select for transformants, G418 (VWR) was added to the medium to 200 mg/L.

### Primers used in this study

The 3’ end sequences bolded in CgSSint1_SS2F and CgSSint1_SS2R anneal to the 5’ and 3’ end regions of the intron respectively. The T boxed in gray is a mismatch designed to modify the 5’ splice site. The fifty 5’ bases match the yeast rDNA flanking the Cas9 cut site. The twenty bases bolded in GuideRNA_SS2R correspond to the guide sequence of plasmid pCAS-SS2-0.

### Plasmid construction by inverse PCR

The guide RNA sequence was designed so the cut site on the yeast rDNA follows the A corresponding to position 534 in the mature SSU yeast rRNA (see Fig. 2). This site was chosen because our mutagenesis plan included the insertion of an intron, and this is where an intron commonly exists in the rDNA of the lichen fungus *Cladonia grayi* (https://genome.jgi.doe.gov/Clagr3/Clagr3.home.html -2018). Due to the position of the PAM sequence, the cut site is one base pair to the left of the exact insertion site of the intron in *C. grayi*. The corresponding guide RNA sequence was inserted into the pCAS plasmid updating the inverse PCR method developed by Hemsley et al. (11) for plasmid mutagenesis. The two primers used, Guide RNA_SS2R and SS2R_ForwExtend (Table 1), have adjacent 5’ ends. Primer Guide RNA_SS2R was designed according to the guidelines in Ryan et al. (25) and contains the desired 20-bp guide RNA sequence flanked on both sides by 20-bp of homologous sequences to the plasmid. The 25 µl reaction contained 15.9 µl dH_2_O, 5 µl Phusion buffer (NEB), 1.25 µl of each 10 µM primer, 0.5 µl dNTPs, 2.5 mM each, 0.125 µl (2ng) pCAS, 1 µl Phusion HF DNA Polymerase (NEB). Thermocycling conditions were: 98°C, 1 min.; 30 cycles of 98°C, 30 sec., 60°C, 1 min., 72°C, 10 min.; 72°C, 10 min. In our hands, inverse PCR worked better than the procedure recommended in Ryan et al. (25) in which two self-complementary primers dramatically reduce yield. With inverse PCR, the desired unmethylated linear product was in large excess relative to the original methylated plasmid, and thus there was no need for DpnI treatment. The linear PCR product was 5’ phosphorylated at 37°C for 30 min in a 20 µl reaction (NEB): 15 µl dH_2_O, 2 µl T4 DNA Ligase buffer, 2 µl PCR product, 1 µl T4 Polynucleotide kinase. For blunt-end ligation, 1 µl T4 DNA Ligase (NEB) was added, and the reaction was incubated at 19°C for 2 hours, producing plasmid pCAS-SS2-0. The latter was introduced into competent *E. coli* strain DH10B and cells were plated on LB-kanamycin medium. Transformants with the correct guide RNA sequence were identified by colony PCR (see below). Plasmid pCAS-SS2-0 was isolated from one of these transformants using a standard alkaline lysis procedure (5) and cleaned with a QIAquick PCR purification kit (Qiagen). The presence of the correct gRNA sequence was confirmed by Sanger sequencing.

**Table 1.**
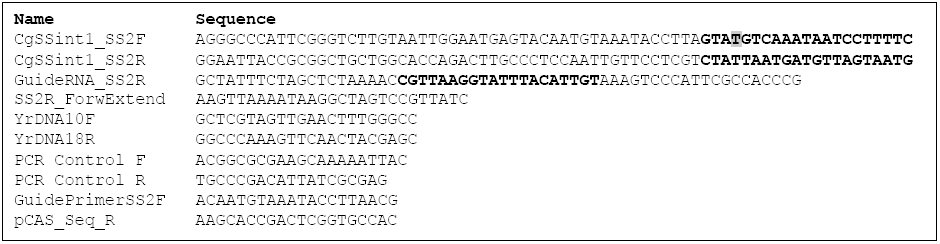

### Intron PCR

The 57-bp intron we chose for transfer into yeast SSU rDNA is from the rDNA of the lichen fungus *Cladonia grayi* (https://genome.jgi.doe.gov/Clagr3/Clagr3.home.html -2018) (Fig. 1). Using *C. grayi* DNA (1) as template, the intron was amplified with two PCR primers, CgSSint1_SS2F and CgSSint1_SS2R (Table 1) whose 21 3‘-end bases primed from the edges of the intron while the 50-bp of their 5’ tails were identical to the yeast rDNA on either side of the Cas9 cut site (Fig. 1 and Table 1). The branchpoint and 3’ splice site sequences of the lichen intron correspond to the most common consensus sequences in yeast (15), but the 5’ splice site corresponds to a less frequent yeast version. To make the 5’ splice site correspond to the most common consensus in yeast, GUAUGU (15), we introduced into the forward primer a one-base A>T mismatch with the original lichen sequence (Fig. 1 and Table 1). The 25 µl PCR contained: 17.3 µl dH_2_O, 5 µl Phusion Buffer (NEB), 0.1 µl of each 100 µM primer, 0.5 µl of dNTPs, 2.5 mM each, 1 µl (5 ng) *Cladonia grayi* DNA, 1 µl Phusion HF DNA Polymerase (NEB). PCR conditions were: 98°C, 30 sec.; 34 cycles of 98°C, 10 sec., 55°C, 30 sec., 72°C, 10 sec.; 72°C, 5 min. The 157-bp fragment obtained was cleaned with a QIAquick PCR purification kit (Qiagen) and used for yeast transformation.

**Figure 1.**
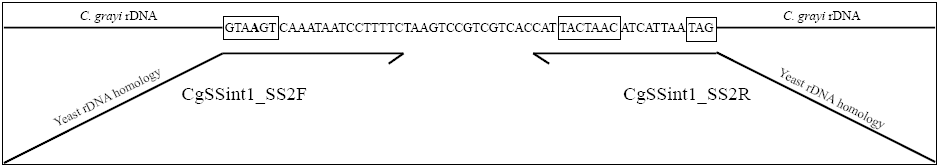
Incorporation of a C. grayi intron into a PCR fragment for integration into yeast rDNA. The top line shows the intron sequence within C. grayi rDNA. From left to right, boxes highlight the 5’ splice site, branchpoint, and 3’ splice site consensus sequences respectively. The bolded A in the 5’ splice site was changed to a T through primer CgSSint1_SS2F. The primer regions annealing to the intron are parallel to the intron. Tilted are the 50-bp regions homologous to yeast rDNA, which allow intron integration into yeast rDNA by HDR. Not drawn to scale.

**Figure 2.**
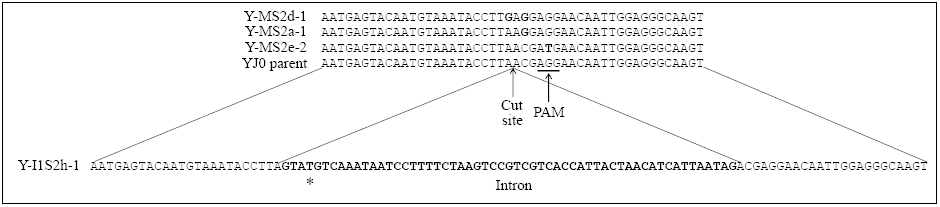
Mutations induced by CRISPR-Cas9 in yeast SSU rRNA around position 534. Shown are the corresponding DNA sequences of four representative transformants. The top is an alignment of three NHEJ mutant sequences and wild type YJ0 around the Cas9 cut site. The substitution mutations are bolded. The bottom sequence is that of a mutant with the C. grayi intron (bolded) inserted at the cut site through HDR. The asterisk corresponds to the base changed through PCR in the 5’ splice site.

### Yeast transformation

The protocol by Ryan et al. (25) was followed for two yeast transformations. One was a co-transformation of 100 µl competent cells with 5 µg intron fragment and 1 µg pCAS-SS2-0 plasmid. Transformants were plated on G418 YEPD plates and incubated at 30°C. The second was a transformation only with pCAS-SS2-0, under the same conditions. Many transformants were slow growing and colonies became visible after 3-4 days. Intron insertion and other mutations were identified by colony PCR (see below) and Sanger sequencing.

### Colony PCR and test of plasmid loss

After introducing plasmid pCAS-SS2-0 into *E. coli* DH10B, transformants were screened for the presence of the correct guide sequence by colony PCR, using one primer specific for the guide sequence, GuidePrimerSS2F, and pCAS_Seq_R as the other primer (Table 1), so that a PCR product would form only in presence of the correct guide sequence. Typically, the 10 µl reaction contained 4.1 µl dH_2_O, 5 µl Apex Taq DNA Polymerase Master Mix solution (Genesee Scientific), 0.2 µl of each 10 µM primer, and 0.5 µl of a diluted *E. coli* suspension. The diluted suspension was obtained by transferring into 10 µl dH_2_O about 0.1 µl of cells picked up with a sterile loop from an *E. coli* colony. PCR conditions were: 94°C, 4 min.; 39 cycles of 94°C, 30 sec., 50°C, 30 sec., 72°C, 15 sec.; 72°C, 7 min.

Colony PCR was performed on yeast transformants to screen for intron insertion and other mutations around the Cas9 cut site. About 0.5-1 µl cells from yeast colonies were disrupted by vortexing and heating for 4 minutes at 90°C in 50 µl TE, 0.1% SDS and 50 µl of 0.3 mm glass beads, and 1 µl of lysate was diluted in 4 µl TE. Typically, the 10 µl reaction contained: 4.4 µl dH_2_O, 5 µl Apex Taq DNA Polymerase Master Mix, 0.2 µl each of 10 µM primer (YrDNA10F and YrDNA18R, Table I), and 0.2 µl of the diluted yeast lysate. Thermocycling conditions were: 95°C, 5 min.; 39 cycles of 95°C, 30 sec., 60°C, 30 sec.; 72°C, 2 min. After prolonged growth in absence of G418, yeast was checked for plasmid loss by colony PCR using plasmid-specific primers PCR Control F and PCR Control R (Table 1). Thermocycling conditions were: 95°C, 5 min.; 39 cycles of 95°C, 30 sec., 60°C, 30 sec.; 72°C, 2 min.

### RNA extraction and cDNA preparation

To demonstrate intron splicing, total RNA was extracted from one of the intron-bearing transformants, Y-I1S2h-1. A 10 ml overnight culture was pelleted in a 50 ml Falcon tube, the pellet was frozen with liquid N_2_ and thoroughly ground in a mortar with liquid N_2_. RNA was extracted using the RNAqueous Total RNA Isolation Kit (ThermoFisher Scientific). The ground cells were resuspended in 1.5 ml of a 0.88:0.22 mix of Lysis solution and Plant RNA isolation aid solution from the kit. Subsequent steps followed the RNAqueous kit protocol. Isolated RNA was quantified by NanoDrop (ThermoFisher Scientific) and quality was assessed by agarose gel electrophoresis. Using Superscript III RT (ThermoFisher Scientific) and the conditions indicated by the manufacturer, reverse transcription was performed at 50°C for 120 min in a 25 µl volume with 100 ng RNA and the reverse primer YrDNA18R (Table 1), whose 5’ end is 111 bases downstream from the intron insertion site. After completion of the reaction, RNA was hydrolyzed with NaOH (final concentration 250 mM). The cDNA was subsequently used as template in PCR using primers YrDNA10F and YrDNA18R (Table 1). PCR conditions were as described for yeast colony PCR. The PCR fragment sizes with and without intron are 290 and 233-bp respectively.

## Results

### CRISPR-Cas9 can effectively mutagenize all nuclear rDNA repeats

Our goal was to test whether CRISPR-Cas9 could be used to homogeneously mutagenize the entire set of 150 rRNA genes in yeast. Plasmid pCAS-SS2-0 was constructed with a guide RNA directing Cas9 to cut the yeast rDNA after the A corresponding to position 534 in the mature SSU rRNA (Fig. 2). We sought to obtain two kinds of mutations at the target site, (i) small base pair changes introduced by non-homologous end joining (NHEJ) and, (ii) the insertion of a 57-bp intron by homology-directed repair (HDR).

Competent yeast was transformed with plasmid pCAS-SS2-0 either by itself or together with a PCR fragment bearing the intron flanked by two 50-bp sequences identical to those spanning the yeast target site. Transformants were screened by colony PCR for presence/absence of intron. Single base pair substitutions around the Cas9 cut site were identified by sequencing. We did not do an exhaustive screen of all mutations occurring around the cut site, but the three substitution patterns shown in Figure 2 recur repeatedly in independent transformants. PCR and sequencing indicate that each change is spread to all repeats: amplifying the intron region with flanking primers in intron-bearing transformants produces only the DNA fragment containing the intron (Fig. 3A); sequence chromatograms of the PCR fragments spanning the small mutations show clean peak patterns, with no indication of heterogeneity (Fig. 3B). All modifications are stably inherited even after removal of the plasmid.

**Figure 3.**
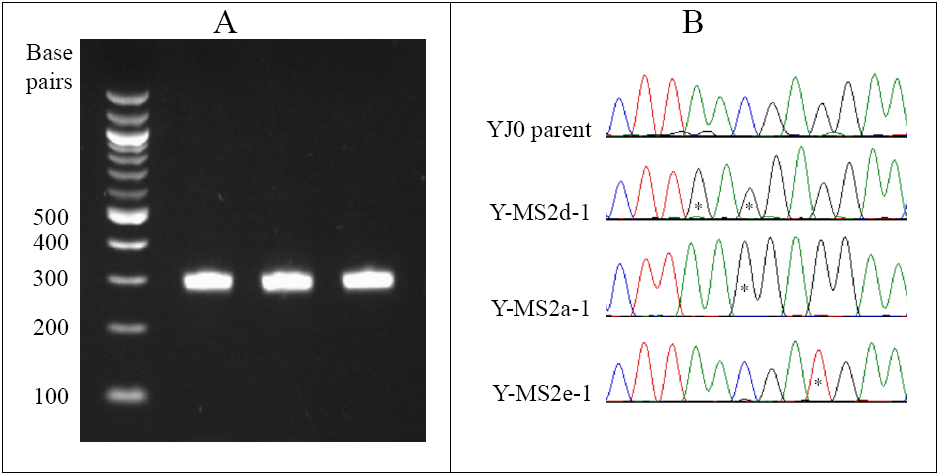
Mutations induced by CRISPR-Cas9 spread to all rDNA repeats. (A) Agarose gel with PCR fragments obtained from three intron mutants using flanking primers YrDNA10F and YrDNA18R. The only visible PCR bands correspond to the 290 bp product containing the intron. The intronless product would be 233 bp long. Size markers are on the left. (B) Sequencing chromatograms of PCR products from the parental YJ0 strain and of three base pair substitution mutants around the Cas9 cut site. The peaks with the substitutions (asterisks) show no minor peaks that would indicate heterogeneity among the amplified rDNA repeats.

### The exogenous intron is correctly spliced by the yeast machinery

The presence of numerous intron-bearing transformants suggested that the yeast was able to correctly splice out the intron during rRNA processing, as its persistence in mature RNA is expected to be lethal. We tested this directly by sequencing a 233-bp cDNA fragment encompassing the intron insertion site, obtained from rRNA extracted from one of the intron-bearing transformants, Y-I1S2h-1. The fragment sequence was identical to the corresponding yeast rRNA sequence, demonstrating precise splicing.

## Discussion

This study is intended as a proof of principle for the use of CRISPR-Cas9 to uniformly spread targeted mutations across all nuclear rDNA repeats in a eukaryote. We counted 647 Cas9 PAM (GGN) sequences within the RDN37-1 rDNA repeat of *S. cerevisiae*, averaging one every 8.2 nucleotides and providing ample potential for intense mutagenesis within this region. The simple method provides a new avenue to study the assembly and structure-function of eukaryotic ribosomes, bypassing the limitations of earlier approaches (see Introduction). The effectiveness of the protocol raises the question of how yeast repairs the concurrent breaks in 150 essential gene copies by Cas9, renders them immune to further cutting by incorporating mutations, and spreads one of those to all repeats quickly enough to survive. The short answer is provided by the convergence of(i) CRISPR’s powerful selection against the persistence of any wild type DNA sequence(ii) a repair system able to contain massive DNA damage (13; 17), and (iii) an rDNA homogenization system spreading the changes to all repeats (28) (9). A detailed answer will have to await experimental dissection of the Cas9 attack on rDNA and of the yeast response to it. The versatility with which yeast can be studied at the molecular level makes it an ideal organism for such analyses. While success in yeast is not a guarantee that the process can be replicated elsewhere, it does increase the likelihood that CRISPR-Cas9 can be effectively used to mutagenize rDNA in other eukaryotes. In one study whose approach resembles ours, CRISPR has been used to inactivate all 62 copies of porcine endogenous retroviruses scattered throughout the pig genome (32).

The three substitutions shown in Figure 2 alter either the PAM sequence or two nucleotides near it, impacting the most critical region for target recognition by Cas9 (26). The mutagenized site in the rRNA is located on SSU helix 17, between bases 534 and 539 (Fig. 4A), on the surface of the small subunit (Fig. 4B). The mutations we introduced fall in two categories. The substitutions resulting from NHEJ of the ds DNA cut persist in the rRNA. They should be either neutral or affect mostly ribosomal structure/function. On the other hand, the intron inserted by HDR does not persist in the rRNA, as survival depends on it being correctly spliced out during rRNA processing. Intron insertion should thus mostly affect rRNA processing and ribosomal assembly. Nuclear rDNA in *S. cerevisiae* and in many other organisms has no introns. However, introns are present in variable numbers in the rDNA of several phyla, including fungi (3). The 57-bp intron used here was obtained from the rDNA of the lichen fungus *Cladonia grayi*. The presence of 5’, branchpoint, and 3’ spliceosomal consensus sequences (Fig. 1), suggests that this intron is spliceosomal. In addition to numerous Group I introns (7), putative spliceosomal introns have been described in the rDNA of lichen fungi (4; 19). Splicing by the yeast machinery indicates that this lichen rDNA intron is indeed spliceosomal. As spliceosomal introns are intimately associated with Polymerase II transcription (12), the ability of lichens and yeast to remove this intron from the Polymerase I rRNA transcript appears to represent an interesting variation of the canonical action of spliceosomes. An additional question is where the normally nucleoplasmic spliceosomes act to remove the intron during ribosomal biogenesis, most of which extends from the nucleolus through the nucleoplasm (31). Although spliceosomes are normally only found in the nucleoplasm (22), there have been sporadic reports suggesting that some mRNA splicing might actually occur in the nucleolus (21). Again, yeast appears to be a good model to address such questions. The biological effects of exogenous rDNA introns in yeast will be discussed elsewhere. It is worth noting here however that rDNA transformants with the intron grow more slowly than both wild type and the rDNA transformants with single base pair mutations. Coping with and splicing out an extraneous intron from rRNA may slow down growth by slowing down assembly and by requiring a significant energy expenditure, not surprising given that ribosomal biogenesis consumes the largest fraction of the cell’s energy (30; 31).

**Figure 4.**
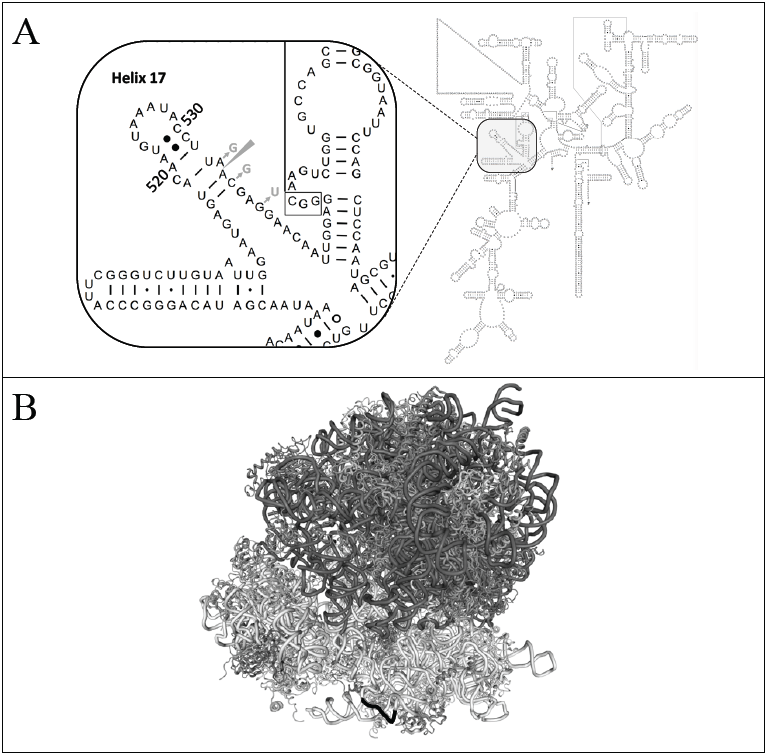
Locations of the induced mutations in the secondary rRNA structure and in the ribosome. (A) The left diagram enlarges the region harboring the mutations and highlighted in the right panel within the secondary structure of the whole S. cerevisiae SSU rRNA (http://www.rna.icmb.utexas.edu/RNA/Structures/d.16.e.S.cerevisiae.pdf - 2017). The mutations, indicated in grey, affect the sequence of helix 17. The wedge points to the location of the Cas9 cut site and of intron insertion in the rDNA. (B) Representation of the full ribosome, with the LSU in dark grey and the SSU in light grey. The mutagenized region of helix 17 is highlighted in black near the bottom of the SSU. Modified from http://www.rcsb.org/pdb/ngl/ngl.do?pdbid=4V7R&bionumber=1 (2017).

